# MolMAE: A Surface-Centric Multimodal Masked Autoencoder for Molecular Representation Learning

**DOI:** 10.64898/2026.07.11.737987

**Authors:** Jiaqing Li

## Abstract

Molecular representation learning has become a central component of modern computational drug discovery. Existing molecular foundation models mainly rely on SMILES strings, two-dimensional molecular graphs, or three-dimensional atomic coordinates. However, many molecular properties are ultimately governed by the molecular surface, where intermolecular recognition, solvation, electrostatic complementarity, and ligand-protein interactions occur. In this work, we propose MolMAE, a surface-guided multimodal masked autoencoder for molecular representation learning. MolMAE takes molecular surface point clouds, three-dimensional molecular graphs, and SMILES-derived fragment and functional-group tokens as complementary input modalities, and learns a unified multimodal molecular embedding through functional-group-aligned masked autoencoding. During pretraining, chemically corresponding local regions are jointly masked across surface, graph, fragment, and functional-group views, forcing the model to reconstruct missing geometric, physicochemical, structural, and semantic information from the remaining context. While molecular surface reconstruction serves as the primary pretraining objective, graph-, fragment-, and functional-group-level reconstruction tasks provide complementary supervision that encourages the model to capture molecular topology, bonding patterns, stereochemistry, local chemical environments, and substructure organization. In addition to reconstructing surface geometry, MolMAE reconstructs surface-associated physicochemical fields, including electrostatic potential and Fukui-related descriptors, enabling the model to learn chemically meaningful surface representations. Pretrained on approximately 261K lead-like bioactive molecules, MolMAE achieves strong performance on the ESOL benchmark under scaffold splitting and competitive performance across multiple molecular property prediction tasks. These results suggest that molecular surface-guided pretraining can complement conventional graph-, sequence-, and atom-coordinate-based molecular representations, especially for property prediction tasks influenced by exposed surface geometry and surface-associated physicochemical patterns.

## 1. Introduction

Molecular property prediction is a fundamental task in computational chemistry and drug discovery, and benchmark suites such as MoleculeNet^1^ have played an important role in standardizing the evaluation of molecular machine learning methods. Recent advances in self-supervised molecular representation learning have led to a variety of molecular foundation models based on SMILES strings, molecular graphs, and three-dimensional conformations. SMILES-based language models, such as ChemBERTa^2^, adapt masked language modeling from natural language processing to molecular strings and learn representations from large-scale unlabeled SMILES corpora. Graph-based models encode atoms and bonds through graph neural networks or graph transformers, with representative self-supervised frameworks including GROVER^3^ and KPGT^4^. Three-dimensional molecular models further incorporate atomic coordinates, distances, and spatial geometry, as demonstrated by geometry-aware pretraining methods such as GEM^5^ and Uni-Mol^6^. Despite these advances, the exposed molecular surface remains underexplored as a pretraining signal, even though it directly mediates interactions with solvents, proteins, membranes, and other molecules.

The molecular surface provides a physically meaningful interface between a molecule and its environment. Classical molecular surface algorithms define solvent-accessible or solvent-excluded surfaces from atomic coordinates, van der Waals radii, and probe radii.^7,8^ More recently, surface-based geometric learning has shown that molecular surfaces encode chemical and geometric fingerprints relevant to biomolecular recognition and protein-ligand interactions.^9^ Many drug-relevant properties are closely related to surface-level physicochemical patterns. Aqueous solubility, for example, is strongly associated with hydrophobicity, molecular size, aromaticity, and conformational flexibility in the ESOL study.^10^ Membrane permeability and oral bioavailability are also related to polar surface area, hydrogen-bonding capacity, and molecular flexibility.^11,12^ In addition, electrostatic potential on the molecular surface reflects charge distribution and long-range electrostatic interactions in solution and biomolecular recognition,^13^ while Fukui-function-based descriptors characterize local chemical reactivity within conceptual density functional theory.^14,15^ These observations suggest that explicitly learning from surface geometry and surface-associated physical fields may provide complementary information beyond graph and SMILES representations.

However, learning from molecular surfaces is challenging. Molecular surfaces are irregular point clouds with molecule-dependent numbers of surface points, and masked modeling on point clouds must handle local patch construction, uneven spatial density, and possible information leakage from point coordinates.^16,17^ Moreover, surface points do not have an explicit one-to-one correspondence with atoms or SMILES tokens. Each local surface patch is generated by nearby atoms and reflects both local molecular geometry and electronic structure. Therefore, an effective surface-based molecular foundation model should not treat the surface as an isolated modality. Instead, it should align surface patches with atomic graph neighborhoods and fragment-level molecular context.

To address these challenges, we introduce MolMAE, a surface-guided multimodal masked autoencoder for molecular representation learning. Inspired by masked language modeling and masked autoencoding,^18,19^ MolMAE takes molecular surface patches, 3D molecular graph neighborhoods, and SMILES-derived fragment and functional-group tokens as complementary molecular views. Unlike conventional molecular masked modeling methods that mainly reconstruct token identities, atom types, graph topology, or atomic coordinates,^3,6,20^ MolMAE performs functional-group-aligned masked pretraining, where chemically corresponding local regions are jointly masked and reconstructed across different views. The surface branch reconstructs local geometry and physicochemical fields, including surface normals, electrostatic potential, and Fukui-related descriptors; the 3D graph branch reconstructs atom- and bond-level structure; and the fragment/functional-group branch reconstructs substructure-level chemical semantics. By aligning these reconstruction tasks within the same masked molecular region, MolMAE learns not only the geometry of the molecular surface, but also the atoms, bonds, and chemical fragments that give rise to surface-level physicochemical patterns. Experiments on molecular property prediction benchmarks show that MolMAE achieves state-of-the-art performance on ESOL after pretraining on approximately 261K lead-like bioactive molecules, while remaining competitive across additional downstream tasks. Ablation and representation analyses further show that molecular surface geometry and surface-associated physicochemical descriptors provide complementary information to graph, fragment, and atom-coordinate-based molecular representations.

## 2. Method

### 2.1 Pretraining Dataset Construction

The pretraining dataset was constructed from the ChEMBL 36 SQLite database^21^ through a multi-stage curation pipeline. Candidate compounds were first restricted to entries annotated as small molecules with valid canonical SMILES. Only parent molecules were retained to remove salt forms, mixtures, and redundant molecular variants. To enrich the dataset with experimentally supported bioactive compounds, we further required each molecule to have at least one standardized activity measurement against a human single-protein target. Activity records were filtered using stringent assay and potency criteria: assay confidence score ≥7, standard activity type in (IC50, Ki, Kd, EC50), and pChEMBL value above the predefined activity threshold.

After bioactivity filtering, molecules were parsed and sanitized using RDKit. Invalid structures, unsupported elements, and duplicated compounds were removed, with deduplication performed by InChIKey. We then applied lead-like physicochemical filters based on molecular weight, calculated LogP, hydrogen-bond donors and acceptors, rotatable bonds, and ring count. This curation procedure yielded 261,840 lead-like, bioactive, non-salt molecules as the initial molecular corpus. For multimodal input construction, each molecule was further processed by 3D conformer generation, NanoShaper^22^-based molecular surface construction, and AIMNet2^23^-based quantum descriptor calculation and surface mapping. Molecules that failed conformer generation, structural validation, surface construction, or descriptor mapping were excluded. After this preprocessing stage, 261,448 molecules were successfully converted into complete surface–graph–fragment multimodal representations and used for MolMAE pretraining.

### 2.2 Molecular Surface Representation

For each molecule, a 3D conformer was prepared and converted into atomic coordinates. The solvent-excluded molecular surface was generated using NanoShaper^22^. The resulting surface was represented as an irregular point cloud containing vertex coordinates, outward surface normals, and precomputed physical descriptors. For a molecule with *N* surface points, the surface representation is defined as 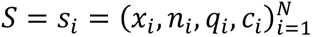, where *x_i_* ∈ ℝ^3^ denotes the coordinate of surface point *i*, *n_i_* ∈ ℝ^3^ is the outward normal vector, *q_i_* ∈ ℝ^3^ denotes QTM-derived quantum/physical descriptors, and *c_i_* ∈ ℝ^3^ denotes multi-scale local curvature descriptors. Functional-group-to-surface associations were further constructed by assigning surface points to nearby functional-group atoms using a distance-based rule. These associations provide the basis for functional-group-aligned surface masking during pretraining. The raw surface point cloud was converted into patch tokens. For each molecule, the number of patch centers was adaptively selected as *G* = clip(round(*N*/8), 64,256).

Patch centers were sampled by farthest point sampling over surface coordinates. For each center, the nearest K=32 surface points were selected to form one patch. The local coordinates in each patch were represented relative to the patch center: *Δx_i,j_*_’_ = *x_i,j_*_’_ – *p_i_*, where *p_i_* is the center of patch *i*. The per-point surface feature was constructed as 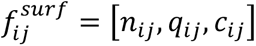. In addition to surface-intrinsic features, each surface point was associated with nearby atoms. The nearest atom types and atom-to-surface distances were collected and processed by a learnable atom-context module. As a result, each surface patch encodes local geometry, surface normals, quantum descriptors, curvature, and nearby atom context.

### 2.3 3D Graph, Fragment, and Functional-Group Representations

In parallel with the molecular surface, each molecule was represented as a 3D molecular graph. Atoms were treated as graph nodes and encoded by atomic number classes. Bonds were treated as graph edges and encoded by bond type together with geometric edge features, including bond length and unit direction vector derived from the 3D conformer. Node chirality labels and a local chirality-geometry scalar were also computed as auxiliary graph targets, providing stereochemical supervision during pretraining.

To introduce higher-level chemical semantics, MolMAE further constructed SMILES-derived fragment tokens and functional-group tokens. Fragment tokens were generated from local atom environments, including atom type, atom degree, chirality label, chirality-geometry bin, and neighboring bond/atom signatures. These fragment signatures were mapped into a fixed-size hashed vocabulary. Functional-group tokens were generated from functional group type, group size, atom composition, and fragment composition. Thus, each molecule was represented by three levels of information: interface-level surface patches, atom-level 3D graph structures, and substructure-level fragment/functional-group tokens. The surface level describes the exposed molecular interface, the graph level provides atom-level topology and 3D geometry, and the substructure level summarizes fragment and functional-group semantics.

The key pretraining strategy of MolMAE is functional-group-aligned co-masking, as shown in Figure 1A. Instead of independently masking surface patches, graph atoms, fragment tokens, and functional-group tokens, MolMAE first selects a valid functional group as the shared masked region. This functional group then determines the masked units across all molecular views. Atoms belonging to the selected functional group are masked in the graph branch, and the graph mask can be further expanded to reach a target atom masking ratio of approximately 20%. Surface patches are masked according to their spatial proximity to the selected functional group. If a precomputed functional-group patch order is available, patches closest to the functional-group center are selected; otherwise, surface points assigned to the functional group are used to seed patch selection. The selected surface patches are constrained to cover approximately 15% of valid patches. Fragment and functional-group masks are derived from the same chemical region: fragment tokens corresponding to masked graph atoms are masked, and the selected functional-group token is masked in the functional-group branch. The masking process can be summarized as

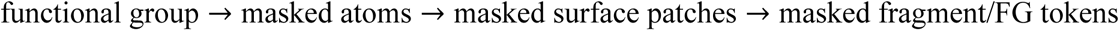

**Figure 1.**
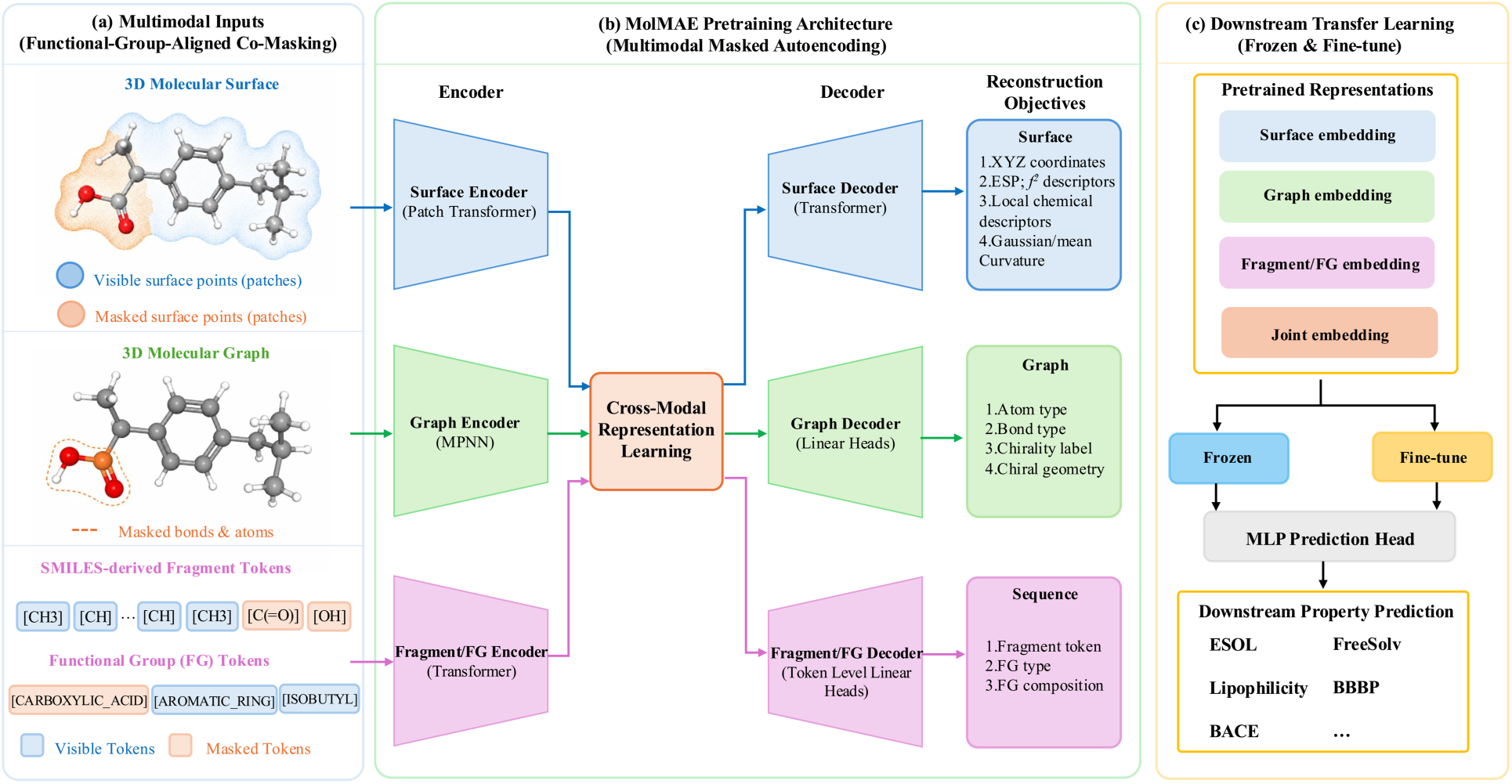
Overview of the MolMAE framework. **(a)** Functional-group-aligned co-masking is performed jointly across three molecular modalities, including the 3D molecular surface, 3D molecular graph, and SMILES-derived fragment/functional-group (FG) tokens, ensuring that semantically corresponding regions are masked simultaneously. **(b)** During pretraining, the three masked modalities are encoded independently and integrated through cross-modal representation learning, comprising surface-graph cross-attention for feature interaction and cross-modal representation alignment to encourage compatible latent representations. **(c)** The pretrained multimodal representations can be either frozen or fine-tuned for downstream molecular property prediction tasks.

This design ensures that the missing information across modalities corresponds to the same local chemical region, encouraging the model to learn relationships among local surface geometry, quantum descriptors, curvature, atom environments, graph topology, fragment signatures, and functional-group semantics.

### 2.4 Model Architecture

MolMAE uses a multimodal masked autoencoding architecture (Figure 1B) consisting of a surface branch, a molecular graph branch, and a fragment/functional-group sequence branch. The surface branch encodes patch tokens with a Transformer encoder. Each surface patch is projected into a hidden dimension of 192, and positional information is added through an MLP applied to the patch center coordinates. Masked surface patches are replaced by a learnable mask token. The encoded surface tokens are pooled to obtain a global surface representation. The graph branch encodes the masked molecular graph and reconstructs atom type, bond type, chirality label, and chirality geometry. By using atom features, bond features, and 3D geometric edge information, the graph branch learns atom-level structural representations that ground the surface representation in molecular topology and stereochemistry. The fragment and functional-group branch uses Transformer encoders over fragment tokens and functional-group tokens. Functional-group representations are further refined using information pooled from corresponding fragment tokens. A joint fusion module combines information from different branches. In the default setting, MolMAE uses surface-graph cross-attention refinement, allowing surface tokens and graph tokens to exchange information directly, while fragment and functional-group tokens provide semantic supervision and cross-modal alignment targets.

### 2.5 Pretraining Objectives

MolMAE is trained with a multimodal masked reconstruction objective. The total loss is defined as

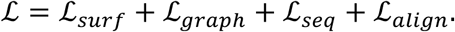

The surface reconstruction loss is

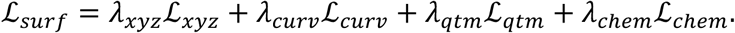

Here, ℒ*_xyz_* reconstructs local surface coordinates, ℒ*_curv_* reconstructs Gaussian curvature and mean curvature, ℒ*_qtm_* reconstructs QTM-derived descriptors, and ℒ*_chem_* reconstructs a distance-weighted local atom-type distribution derived from nearby atoms. This objective encourages the surface branch to learn both local molecular shape and chemically meaningful surface fields. The graph reconstruction loss is

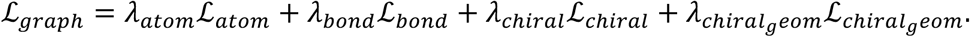

Atom type, bond type, and chirality label are optimized using cross-entropy loss, while chirality geometry is optimized using Smooth L1 loss. This objective allows the graph branch to recover atom identity, bond topology, 3D geometric structure, and stereochemical information from masked molecular regions. The sequence-level reconstruction loss is

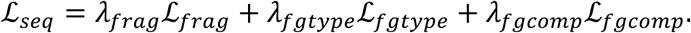

Here, ℒ*_frag_*, reconstructs masked fragment tokens, ℒ*_f gtype_* predicts functional-group type, and ℒ*_f gcomp_*. reconstructs functional-group composition. These objectives provide substructure-level semantic supervision and help the model learn chemically meaningful fragment and functional-group representations. In addition, MolMAE uses Smooth L1 alignment losses to connect corresponding representations across modalities:

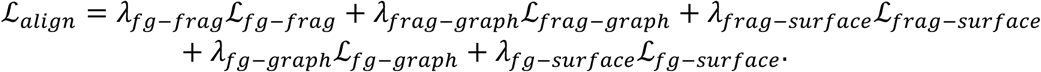

These alignment losses encourage functional-group, fragment, graph, and surface representations associated with the same chemical region to occupy compatible latent spaces.

To stabilize multimodal pretraining, MolMAE adopts a staged optimization schedule in which different reconstruction and alignment objectives are activated progressively. The training starts with the most fundamental reconstruction tasks, including surface coordinate reconstruction and graph atom/bond reconstruction. These objectives provide stable geometric and topological supervision and allow the model to first learn local surface structure and molecular graph connectivity.

### 2.6 Pretraining Schedule and Optimization Focus

After the initial stage, surface-level physical reconstruction objectives are introduced, including QTM descriptor reconstruction and local chemical atom-distribution reconstruction. These losses encourage the surface branch to move beyond pure geometric reconstruction and learn physically meaningful surface fields and nearby atom environments. In the graph branch, chirality-related supervision is activated after the model has learned stable atom and bond representations. Specifically, MolMAE reconstructs both discrete chirality labels and a continuous chirality-geometry scalar, which provides additional stereochemical information for molecular representation learning. Higher-level semantic objectives are introduced in later stages. Fragment reconstruction is used to supervise local chemical environment tokens, while functional-group type and composition prediction provide substructure-level chemical supervision. Finally, cross-modal alignment losses are activated to align functional-group, fragment, graph, and surface representations associated with the same masked chemical region. This delayed activation prevents early training from being dominated by noisy semantic or alignment targets before the geometry and graph encoders become stable. This staged strategy reflects the design priority of MolMAE: first learning reliable local geometry and atom-level structure, then incorporating surface physical fields, stereochemistry, fragment semantics, functional-group information, and finally cross-modal consistency.

### 2.7 Transfer Learning for Downstream Tasks

After pretraining, the learned molecular representations are transferred to downstream molecular property prediction tasks through a transfer learning paradigm. Specifically, the pretrained MolMAE encoder serves as a general-purpose molecular feature extractor and is subsequently fine-tuned on task-specific datasets. This strategy enables knowledge acquired from large-scale self-supervised pretraining, including molecular surface geometry, physicochemical surface fields, graph topology, stereochemistry, fragment semantics, and functional-group information, to be effectively adapted to diverse prediction tasks with limited labeled data. The transferred representations are evaluated on standard molecular property benchmarks, including regression tasks such as ESOL, FreeSolv, and Lipophilicity, as well as classification tasks such as BBBP and BACE.

## 3. Experiments and Results

To evaluate the effectiveness of MolMAE, we conducted experiments on multiple molecular property prediction tasks. Compared with conventional sparse graph- or string-based molecular representations, MolMAE operates on dense molecular surface point clouds enriched with quantum-derived physical descriptors, which increases the computational cost of pretraining and downstream fine-tuning. Accordingly, we focus our transfer evaluation on small- to medium-scale molecular property prediction benchmarks, where labeled data are often limited and the quality of pretrained molecular representations is particularly important. This evaluation setting allows us to assess the practical transferability of MolMAE under realistic downstream conditions while maintaining a balance between computational cost and benchmark coverage. The results demonstrate that surface-guided multimodal pretraining can provide effective molecular representations for downstream property prediction despite the additional cost introduced by surface-based modeling.

### 3.1 Molecular Property Prediction Setup

We evaluated MolMAE on seven molecular property prediction datasets from MoleculeNet^1^, including three regression tasks, ESOL, FreeSolv, and Lipophilicity, and four classification tasks, BBBP, BACE, ClinTox, and SIDER. Following Uni-Mol^6^, we used scaffold splitting to evaluate model generalization to structurally distinct molecules. All results are reported over three random seeds, 0, 1, and 42. For a fair comparison, all MolMAE variants and baselines are evaluated using exactly matched scaffold split indices and the same downstream evaluation protocol.

For regression tasks, we reported root mean squared error (RMSE), where lower values indicate better performance. For classification tasks, we reported ROC-AUC in percentage, where higher values indicate better performance. For multi-task classification datasets such as ClinTox and SIDER, macro-averaged ROC-AUC was used. The number of molecules reported in the tables refers to the valid molecules retained after preprocessing and downstream data validation.

We compared MolMAE against Uni-Mol as the primary reference baseline, because Uni-Mol is a strong atom-coordinate-based 3D molecular representation model that has been extensively evaluated against multiple competitive baselines on MoleculeNet benchmarks and was reported to achieve state-of-the-art or near state-of-the-art performance across most evaluated tasks. In addition to Uni-Mol, we included two MolMAE ablation variants to analyze the contribution of molecular surface information. MolMAE-NoSurf removes the molecular surface branch and therefore evaluates the contribution of graph, fragment, and functional-group information without explicit surface modeling. MolMAE-SurfGeo retains the 3D surface branch but uses only surface geometric information, including surface coordinates, local geometry, curvature, and nearby atom context, while removing surface QTM descriptors and local chemical-environment reconstruction. The full MolMAE further incorporates surface quantum/physical descriptors and local chemical-environment reconstruction.

### 3.2 Downstream Fine-tuning Performance

Results are summarized in Table 1 and Table 2, where the best performance for each benchmark is highlighted in bold. Table 1 reports regression performance on ESOL, FreeSolv, and Lipophilicity, while Table 2 reports classification performance on BBBP, BACE, ClinTox, and SIDER.

**Table 1.** MolMAE performance on molecular property prediction regression tasks.

| Regression (RMSE, lower is better ↓) |  |  |  |  |
| --- | --- | --- | --- | --- |
| Datasets | ESOL | FreeSolv | Lipophilicity |  |
| #Molecules | 1128 | 641 | 4,200 |  |
| #Tasks | 1 | 1 | 1 |  |
| Uni-Mol | $0.867 \pm 0.156$ | **$0.836 \pm 0.177$** | **$0.644 \pm 0.068$** | |
| MolMAE-SurfGeo | <u>$0.773 \pm 0.081$</u> | $1.086 \pm 0.403$ | <u>$0.700 \pm 0.057$</u> | |
| MolMAE-NoSurf | $0.785 \pm 0.082$ | $1.052 \pm 0.317$ | $0.733 \pm 0.065$ | |
| MolMAE | **$0.740 \pm 0.091$** | <u>$0.967 \pm 0.248$</u> | $0.710 \pm 0.063$ | |

**Table 2.** MolMAE performance on molecular property prediction classification tasks Classification (ROC-AUC (%), higher is better ↑)

| Classification (ROC-AUC (%), higher is better ↑) |  |  |  |  |
| --- | --- | --- | --- | --- |
| Datasets | BBBP | BACE | ClinTox | SIDER |
| #Molecules | 1,994 | 1,513 | 1,421 | 1,137 |
| #Tasks | 1 | 1 | 2 | 27 |
| Uni-Mol | $85.05 \pm 2.24$ | $81.48 \pm 12.50$ | $76.35 \pm 7.85$ | <u>$59.22 \pm 4.06$</u> |
| MolMAE-SurfGeo | $91.47 \pm 1.17$ | $81.80 \pm 6.20$ | **$77.87 \pm 8.74$** | $58.66 \pm 4.88$ |
| MolMAE-NoSurf | <u>$91.82 \pm 2.62$</u> | <u>$82.88 \pm 8.24$</u> | $65.72 \pm 5.42$ | $58.53 \pm 3.22$ |
| MolMAE | **$93.72 \pm 1.34$** | **$85.86 \pm 6.29$** | <u>$77.15 \pm 7.61$</u> | **$59.45 \pm 2.00$** |

On regression tasks, MolMAE achieves the best performance on ESOL, reducing the RMSE from 0.867 for Uni-Mol to 0.740. The ablation results further support the contribution of molecular surface information. MolMAE-NoSurf obtains an RMSE of 0.785, while MolMAE-SurfGeo improves it to 0.773, indicating that surface geometry alone already provides useful information for solubility prediction. The full MolMAE further improves the RMSE to 0.740, suggesting that surface-associated quantum/physical descriptors and local chemical-environment reconstruction provide additional benefits beyond geometry-only surface modeling. This trend is chemically reasonable because aqueous solubility is closely related to exposed molecular shape, surface polarity, local electrostatics, and solute-solvent interactions.

Because ESOL is the benchmark where MolMAE shows the strongest improvement, we further compare MolMAE with additional supervised graph-based baselines commonly used for aqueous solubility prediction. Previous solubility studies have identified Chemprop^24^ and AttentiveFP^25^ as strong deep learning baselines for ESOL and related solubility datasets.^26^ To avoid confounding effects from different data splits or evaluation workflows, we reported all baselines under the same scaffold-split protocol used in this study. Chemprop is reported using the best setting selected from the grid-search protocol, while Uni-Mol is evaluated using its official recommended setting. As shown in Table S1, MolMAE achieves the lowest average RMSE on ESOL, slightly outperforming Chemprop and AttentiveFP and clearly outperforming Uni-Mol under this evaluation setting. MolMAE consistently achieves the lowest RMSE under the matched scaffold-split protocol, demonstrating the effectiveness of surface-guided multimodal pretraining for ESOL prediction.

A similar trend is observed on FreeSolv, where the full MolMAE improves over both MolMAE-NoSurf and MolMAE-SurfGeo, although Uni-Mol remains stronger overall. The improvement from MolMAE-SurfGeo to the full MolMAE indicates that surface physicochemical reconstruction contributes additional information, but the current single-conformer surface representation may still be insufficient for properties that depend strongly on conformational ensembles and thermodynamic effects. On Lipophilicity, MolMAE-SurfGeo slightly outperforms the full MolMAE, suggesting that the benefit of surface physicochemical reconstruction is task dependent. Lipophilicity may depend more strongly on global hydrophobic balance, conformational variability, and dataset-specific calibration than on a single-conformer molecular surface alone.

On classification tasks, MolMAE shows strong performance on BBBP and BACE. On BBBP, MolMAE achieves a ROC-AUC of 93.72%, outperforming Uni-Mol by 8.67 percentage points. Both MolMAE-NoSurf and MolMAE-SurfGeo also outperform Uni-Mol on this dataset, suggesting that the graph, fragment, and functional-group components of the model already learn useful transferable features for permeability-related prediction. The full MolMAE further achieves the highest performance, indicating that surface physicochemical information provides an additional improvement. On BACE, the full MolMAE also achieves the best ROC-AUC of 85.86%, outperforming Uni-Mol and both ablation variants. Together, these observations show that combining surface geometry, surface physical descriptors, atom-level graph structure, and fragment-level chemical semantics is beneficial for bioactivity-related classification.

The ablation results indicate that MolMAE-NoSurf, MolMAE-SurfGeo, and the full MolMAE should not be interpreted as a strictly monotonic sequence of increasingly stronger models. Instead, they represent different inductive biases. MolMAE-NoSurf emphasizes graph topology, fragment tokens, and functional-group semantics. MolMAE-SurfGeo emphasizes 3D surface geometry, curvature, and nearby atom context without explicit surface physicochemical reconstruction. The full MolMAE combines surface geometry with QTM-derived descriptors and local chemical-environment reconstruction.

This task-dependent behavior is most evident on ClinTox. MolMAE-SurfGeo slightly outperforms the full MolMAE on ClinTox, but the difference is small relative to the standard deviation and should not be overinterpreted. ClinTox is a small multi-task toxicity dataset with substantial label imbalance, sparse supervision, and high split sensitivity. In this setting, the larger full model and additional surface physicochemical objectives may introduce extra fine-tuning variance or mild overfitting, whereas the simpler geometry-only surface variant may act as a more regularized representation. Therefore, the slightly better ClinTox performance of MolMAE-SurfGeo should be interpreted as a task-dependent variance effect rather than evidence that surface physicochemical descriptors are generally unhelpful.

On SIDER, the performance differences among Uni-Mol, MolMAE-SurfGeo, MolMAE-NoSurf, and the full MolMAE are relatively small. The full MolMAE obtains the best average ROC-AUC, but the margin is modest. This suggests that side-effect prediction may not be determined by local molecular surface patterns alone and may require additional biological context, target/pathway information, or larger-scale supervised data. Overall, the results show that MolMAE provides clear benefits on selected downstream tasks, especially ESOL, BBBP, and BACE, while remaining complementary rather than universally superior to Uni-Mol across all benchmarks.

### 3.3 Linear-Probe Evaluation

To further evaluate the intrinsic quality of the pretrained representations, we conducted a linear-probe experiment on the same seven downstream benchmarks. In this setting, the pretrained encoder is frozen and only a lightweight prediction head is trained. Therefore, the linear-probe results reflect how much task-relevant information is already encoded in the learned molecular embeddings before full end-to-end fine-tuning.

Table 3 reports the linear-probe performance of MolMAE, Uni-Mol, MolMAE-SurfGeo, and MolMAE-NoSurf on all seven downstream tasks. MolMAE achieves the best linear-probe performance on all three regression tasks, including ESOL, FreeSolv, and Lipophilicity. The superior linear-probe performance indicates that the full surface-guided multimodal encoder captures strong physicochemical information in its frozen representation. In particular, the improvement over Uni-Mol on these regression benchmarks indicates that molecular surface geometry, QTM-derived descriptors, and local chemical-environment reconstruction provide useful information beyond atom-coordinate-based 3D pretraining.

**Table 3.** Linear-probe performance on seven molecular property prediction tasks. (Results are reported as mean ± standard deviation over matched scaffold-split seeds. For regression tasks, lower RMSE is better. For classification tasks, higher ROC-AUC (%) is better. Bold values indicate the best result for each task.)

| Task | MolMAE | Uni-Mol | MolMAE-SurfGeo | MolMAE-NoSurf |
| --- | --- | --- | --- | --- |
| ESOL | **0.9221 ± 0.0576** | <u>1.0257 ± 0.0829</u> | 1.0357 ± 0.0661 | 1.0761 ± 0.0430 |
| FreeSolv | **1.4845 ± 0.1414** | <u>1.5979 ± 0.0495</u> | 1.6924 ± 0.2152 | 1.6027 ± 0.4486 |
| Lipophilicity | **0.8466 ± 0.0495** | 1.0092 ± 0.1008 | 0.8973 ± 0.1057 | <u>0.8845 ± 0.1013</u> |
| BBBP | 83.89 ± 2.96 | 82.83 ± 4.61 | <u>84.11 ± 5.01</u> | **86.12 ± 3.48** |
| BACE | <u>81.34 ± 7.51</u> | 77.76 ± 8.01 | 80.79 ± 6.51 | **82.87 ± 7.06** |
| ClinTox | 78.18 ± 7.41 | **83.34 ± 6.45** | <u>80.07 ± 4.73</u> | 70.87 ± 2.74 |
| SIDER | <u>58.17 ± 3.45</u> | 57.07 ± 1.09 | 55.19 ± 3.59 | **58.88 ± 2.15** |

For classification tasks, the linear-probe results are more task dependent. MolMAE-NoSurf obtains the best linear-probe performance on BBBP, BACE, and SIDER, while Uni-Mol performs best on ClinTox. This suggests that graph, fragment, and functional-group semantics can already provide strong linearly accessible signals for several classification tasks, especially when the encoder is frozen. Nevertheless, the full MolMAE remains competitive across all classification benchmarks and becomes stronger after end-to-end fine-tuning on BBBP and BACE. Together, the linear-probe and fine-tuning results indicate that MolMAE learns transferable representations, while different downstream tasks benefit from different balances of surface, graph, fragment, and functional-group information.

### 3.4 Representation-Level Complementarity Between MolMAE and Uni-Mol

To examine whether MolMAE provides information beyond Uni-Mol rather than simply reproducing an existing 3D molecular representation, we compared their embedding spaces using centered kernel alignment and nearest-neighbor overlap. As shown in Figure 2, MolMAE shows only moderate similarity to Uni-Mol, with an overall CKA of 0.44 and an Overlap@10 of 0.32. This indicates that the two models organize molecules differently in representation space.

**Figure 2.**
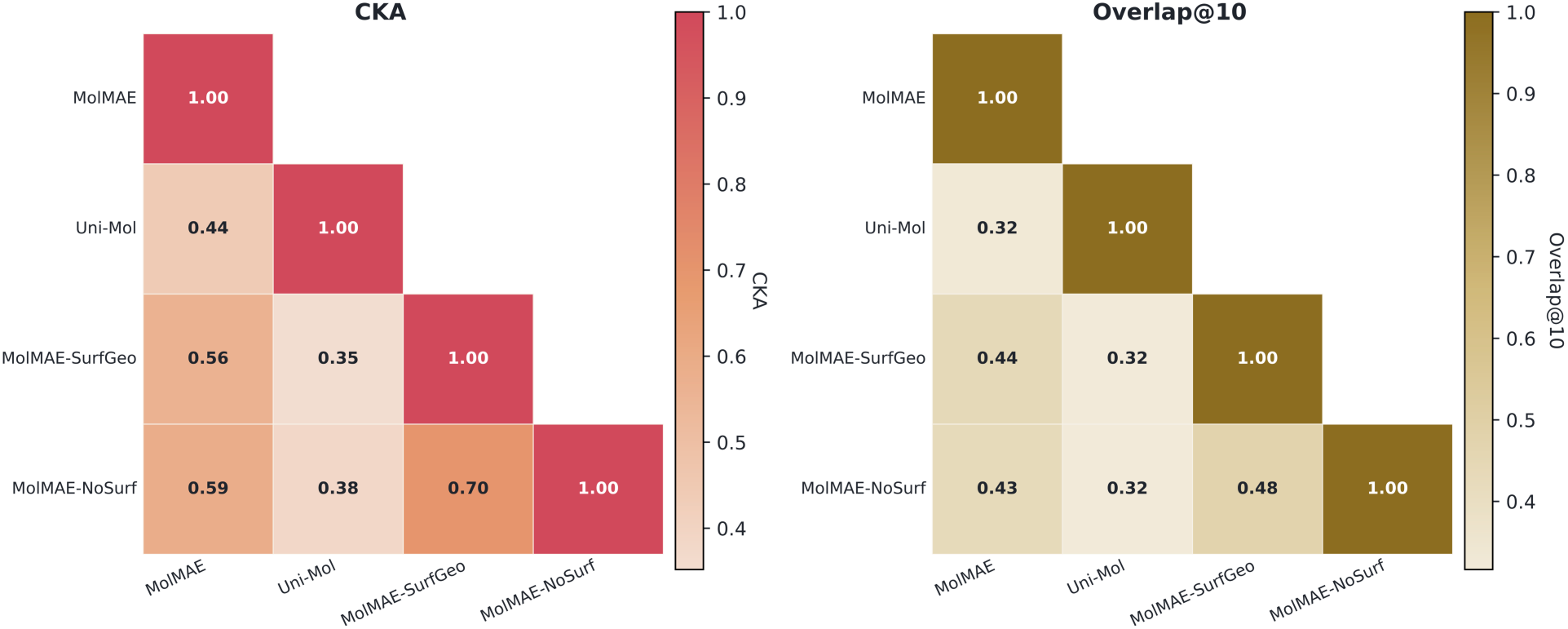
Overall representation similarity among MolMAE, Uni-Mol, and surface ablation variants.

In contrast, MolMAE is more closely related to its own ablation variants than to Uni-Mol, as expected from their shared graph, fragment, and functional-group components. MolMAE-SurfGeo and MolMAE-NoSurf also show relatively high similarity to each other, suggesting that geometry-only surface modeling remains closer to the non-surface representation than the full surface-physicochemical representation. These results support the interpretation that full MolMAE introduces a distinct surface-guided inductive bias through QTM-derived descriptors and local chemical-environment reconstruction.

This representation-level analysis is important because downstream performance alone cannot determine whether MolMAE learns genuinely new information or merely duplicates existing 3D molecular embeddings. The moderate similarity between MolMAE and Uni-Mol suggests that molecular surface pretraining captures complementary information to atom-coordinate-based 3D pretraining. Therefore, MolMAE and Uni-Mol should be viewed as complementary molecular representation models rather than mutually exclusive alternatives. Additional task-wise CKA and Overlap@10 matrices are provided in Figure S1, showing that the representation-level relationship among MolMAE, Uni-Mol, and the ablation variants varies across downstream datasets. Low-dimensional embedding visualizations in Figure S2 further show that the four encoders organize the same molecular collections in different ways. We also evaluate predictive complementarity by concatenating MolMAE and Uni-Mol embeddings in a linear-probe setting; the corresponding gain analysis is shown in Figure S4. Positive gains from embedding combination provide additional evidence that surface-guided and atom-coordinate-based representations contain non-redundant molecular information.

### 3.5 Qualitative Surface Reconstruction and Property Recovery

We further qualitatively evaluate whether MolMAE learns meaningful surface representations through masked surface reconstruction. Figure 3 shows a representative validation molecule, where the model is given a masked surface input and asked to reconstruct both the missing surface region and surface-associated scalar fields.

**Figure 3.**
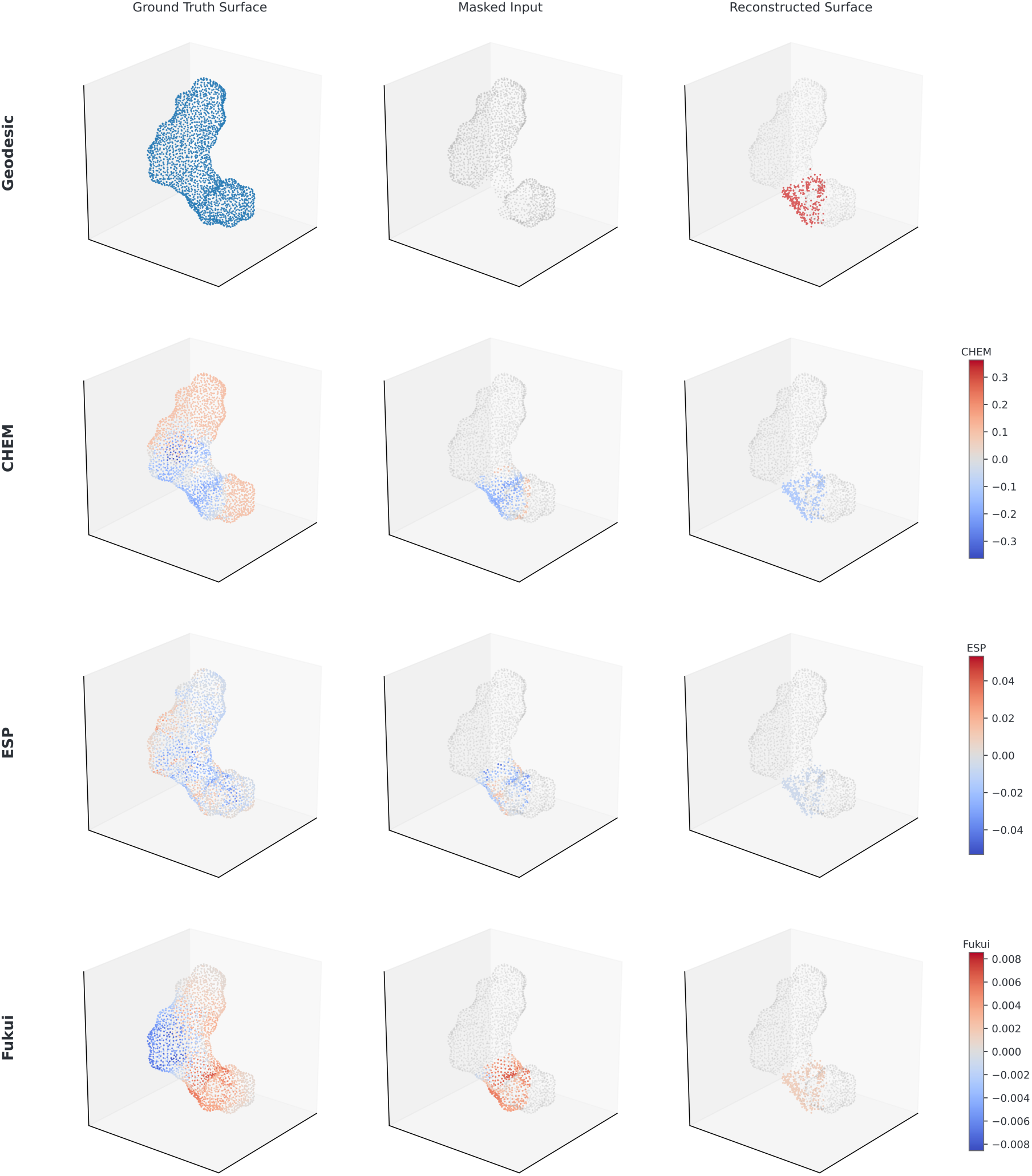
Qualitative masked-surface reconstruction and surface-property recovery on a representative validation molecule (CHEMBL1332397). A representative validation molecule is shown to illustrate MolMAE’s ability to reconstruct masked molecular surface regions and associated surface fields. Columns correspond to the ground-truth surface, masked input, and reconstructed surface, respectively. Rows show the reconstructed targets at different levels: surface geometry, local chemistry (CHEM), ESP, and Fukui descriptors. In the masked input, the visible surface is shown in gray and the masked region is colored by the corresponding ground-truth scalar field. The reconstructed surface panel shows the model prediction for the masked region overlaid on the visible context. MolMAE recovers the local geometry of the missing surface patch and preserves the major spatial patterns of CHEM, ESP, and Fukui fields, although the predicted scalar fields are smoother than the ground-truth distributions.

The reconstructed surface recovers the masked local region, and the predicted local chemical-environment field (CHEM), ESP, and Fukui-related fields preserve the major spatial patterns of the corresponding ground-truth distributions. Although the reconstructed scalar fields are smoother than the ground truth, they capture the local sign and spatial organization of the masked region. This qualitative result suggests that MolMAE learns more than geometric point completion: it also captures chemically meaningful surface properties associated with the masked molecular region.

This observation supports the design of the full pretraining objective. A geometry-only masked autoencoder may learn to infer missing coordinates from neighboring surface points, but it does not necessarily learn the physicochemical meaning of the reconstructed region. By jointly reconstructing surface geometry, QTM-derived descriptors, and local chemical-environment targets, MolMAE is encouraged to encode both the shape and the chemical identity of molecular surfaces.

## 4. Discussion

This study investigates whether explicit molecular surface modeling can provide useful information beyond conventional graph-, fragment-, and atom-coordinate-based molecular representations. Our experiments show that the answer is task dependent but generally positive. Compared with Uni-Mol, MolMAE achieves stronger performance on ESOL, BBBP, and BACE, while Uni-Mol remains stronger on FreeSolv, Lipophilicity, and ClinTox. Therefore, MolMAE should not be interpreted as a universal replacement for Uni-Mol. Instead, it should be viewed as a surface-guided multimodal representation that captures complementary molecular information.

The strongest evidence for the value of molecular surface modeling comes from ESOL. MolMAE outperforms Uni-Mol and both ablation variants, and performance improves from MolMAE-NoSurf to MolMAE-SurfGeo and further to the full MolMAE. The progressive improvement indicates that surface geometry and surface-associated physicochemical descriptors contribute complementary information for solubility prediction. This is chemically reasonable because aqueous solubility depends strongly on exposed molecular shape, local polarity, electrostatic patterns, and solute-solvent interactions. A molecular graph or atom-coordinate representation can encode structural information, but it does not explicitly represent the exposed molecular interface. By modeling molecular surfaces directly, MolMAE provides a more interface-centered view of the molecule.

The ablation results clarify why the full MolMAE model is preferred. MolMAE-NoSurf captures graph, fragment, and functional-group semantics without explicit surface modeling. MolMAE-SurfGeo adds surface geometry, curvature, and nearby atom context, but does not use surface QTM descriptors or local chemical-environment reconstruction. The full MolMAE combines surface geometry with physicochemical surface supervision. The performance gains observed for the full model on several tasks suggest that reconstructing surface coordinates alone may be insufficient. For molecular surfaces, two local regions can have similar shape but very different electrostatic or chemical properties. Therefore, reconstructing QTM-derived descriptors and local chemical environments forces the model to learn not only what the surface looks like, but also what the surface means chemically.

At the same time, the ablation results show that the three MolMAE variants are not simply a monotonic sequence of increasingly better models. They represent different inductive biases. MolMAE-NoSurf emphasizes chemical topology and substructure semantics. MolMAE-SurfGeo emphasizes surface geometry with a relatively simpler supervision signal. The full MolMAE adds additional physicochemical reconstruction objectives and therefore has greater expressive power but also greater fine-tuning complexity. This explains why MolMAE-SurfGeo can slightly outperform the full model on ClinTox. ClinTox is a small multi-task toxicity benchmark with high variance, label imbalance, and limited supervision. In such settings, the simpler geometry-only variant may act as a stronger regularized representation, while the full model may introduce additional fine-tuning variance. Because the difference is small relative to the standard deviation, this result should be interpreted as a task-dependent variance effect rather than evidence against surface physicochemical supervision.

The comparison with Uni-Mol provides an important perspective on the role of molecular surfaces. Uni-Mol is a strong atom-coordinate-based 3D representation model and remains better on some physicochemical tasks. One likely reason is that the current MolMAE uses a single conformer and its corresponding molecular surface, while properties such as hydration free energy and lipophilicity can depend on conformational ensembles, global thermodynamic effects, and dataset-specific calibration. A single surface representation may not fully capture these effects. Incorporating multi-conformer molecular surfaces or conformer-level pooling may therefore improve future versions of MolMAE. In addition, the molecular-weight distributions of the MolMAE pretraining corpus and the downstream benchmarks differ across datasets, as shown in Figure S3, which may partially contribute to the task-dependent transfer behavior observed in our experiments.

The representation-level analyses support the interpretation that MolMAE and Uni-Mol are complementary. Under linear probing, MolMAE provides strong frozen representations for regression tasks, suggesting that its pretrained embedding already contains useful physicochemical information. The CKA and nearest-neighbor overlap analyses further show that MolMAE and Uni-Mol organize molecules differently in embedding space. This is important because performance alone cannot distinguish whether MolMAE learns genuinely new information or merely reproduces existing 3D molecular representations. The moderate similarity between MolMAE and Uni-Mol indicates that explicit surface-guided pretraining introduces a distinct inductive bias.

The qualitative reconstruction analysis provides additional evidence that the model learns chemically meaningful surface information. MolMAE is able to reconstruct masked surface regions and recover associated scalar fields such as CHEM, ESP, and Fukui descriptors. This supports the design of the full pretraining objective, where geometry reconstruction is combined with physicochemical reconstruction. In this sense, MolMAE differs from a purely geometric point-cloud autoencoder: the goal is not only to complete missing surface coordinates, but also to learn the physicochemical identity of the exposed molecular interface.

Overall, these results suggest that molecular surfaces provide an informative and complementary representation for molecular property prediction, especially for tasks where exposed surface geometry, local physicochemical fields, and functional groups are directly related to the target property, such as ESOL, BBBP, and BACE. For tasks that depend more strongly on conformational ensembles, biological context, or sparse multi-task labels, the current model remains limited. In addition, surface-based modeling introduces higher computational cost than atom-coordinate-based approaches such as Uni-Mol, because MolMAE requires molecular surface generation, descriptor mapping, patch construction, and dense point-cloud processing. These steps increase preprocessing time and memory consumption, indicating that the efficient and interpretable extraction of molecular surface information remains an important challenge. Future work should focus on multi-conformer surface pretraining, improved regularization for small datasets, fusion strategies that combine MolMAE and Uni-Mol embeddings, faster and less redundant surface sampling, and more interpretable methods for identifying task-relevant surface descriptors and learned surface patches.

## Supplementary Materials

To support the representation-level interpretation in the main text, we provide additional analyses examining task-wise embedding similarity, chemical-space organization, dataset distribution, and predictive complementarity between MolMAE and Uni-Mol.

Task-wise representation similarity is reported in Figure S1 using centered kernel alignment (CKA) and nearest-neighbor Overlap@10 among MolMAE, Uni-Mol, MolMAE-SurfGeo, and MolMAE-NoSurf. The similarity patterns vary across downstream datasets, indicating that the relationships among different representations are task dependent. Across several tasks, MolMAE is more similar to its ablation variants than to Uni-Mol, suggesting that MolMAE learns a representation space that is related to, yet distinct from, atom-coordinate-based 3D pretraining. This analysis complements the overall representation similarity results presented in the main text.

Low-dimensional projections of the learned embeddings are shown in Figure S2. These visualizations provide a qualitative view of how MolMAE, Uni-Mol, MolMAE-SurfGeo, and MolMAE-NoSurf organize the same molecular collections in representation space. The different spatial organizations observed across models are consistent with the CKA and Overlap@10 analyses, suggesting that molecular surface pretraining leads to a distinct organization of chemical space rather than simply reproducing Uni-Mol embeddings.

Figure S3 compares the molecular-weight distributions of the MolMAE pretraining corpus and the seven downstream benchmarks. The 261K MolMAE pretraining corpus is concentrated within a lead-like molecular-weight regime, whereas the downstream datasets exhibit distinct and, in some cases, broader molecular-weight distributions. This distributional discrepancy provides useful context for interpreting the task-dependent transfer behavior observed in the main results.

Predictive complementarity between MolMAE and Uni-Mol embeddings is examined in Figure S4. We concatenate the two embeddings and compare the resulting linear-probe performance with that obtained from each single-model embedding. Improvements from embedding combination indicate that surface-guided and atom-coordinate-based representations capture non-redundant molecular information, providing additional evidence that MolMAE and Uni-Mol encode complementary molecular features.

**Figure S1.**
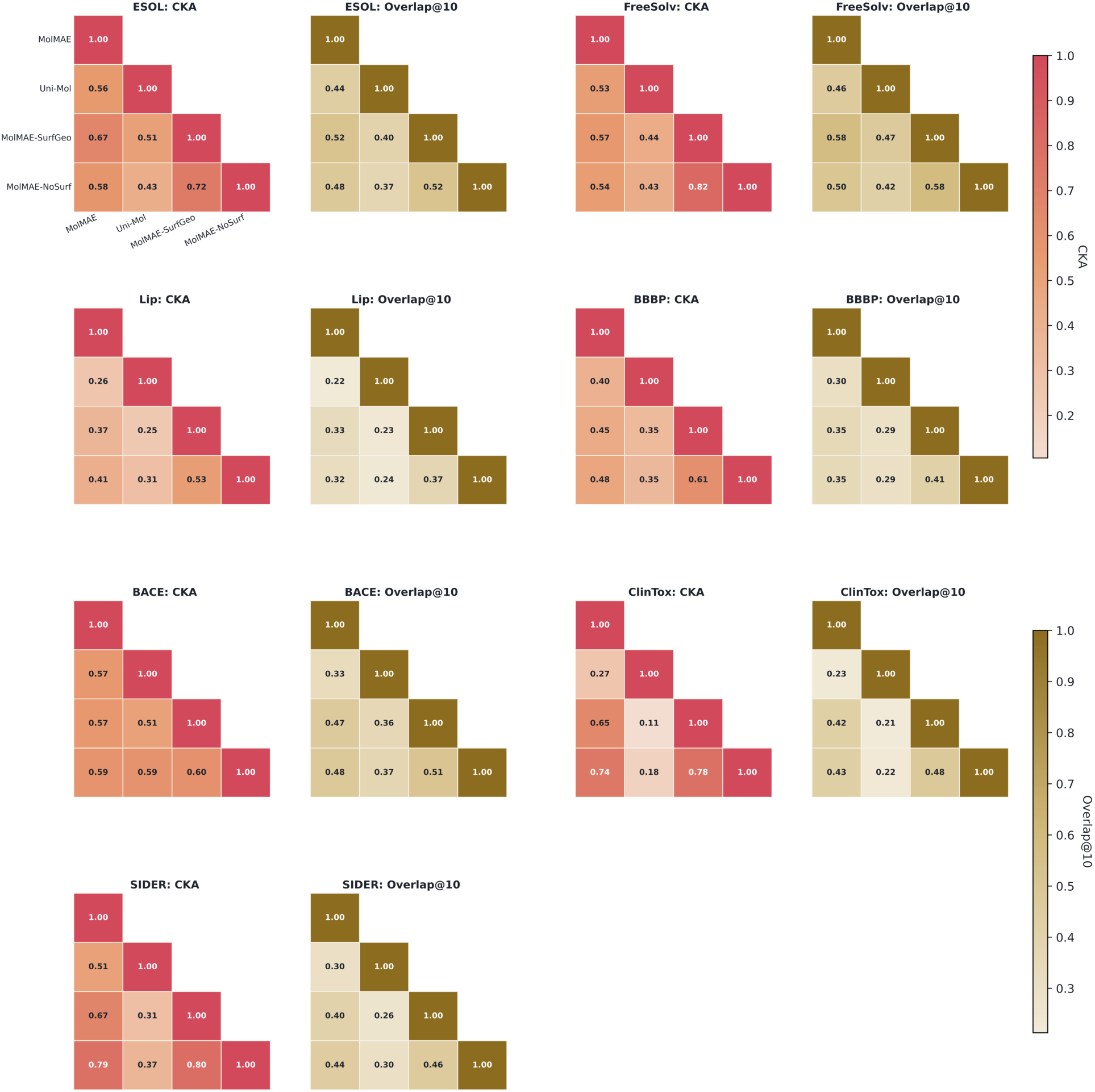
Task-wise representation similarity across seven molecular property prediction benchmarks.

Pairwise representation similarity among MolMAE, Uni-Mol, MolMAE-SurfGeo, and MolMAE-NoSurf is computed separately for each downstream task using centered kernel alignment (CKA) and nearest-neighbor Overlap@10. The heatmaps show that representation similarity is task dependent. MolMAE is generally more closely related to its ablation variants than to Uni-Mol, while the MolMAE-Uni-Mol similarity remains moderate across tasks. This supports the interpretation that surface-guided pretraining induces a representation space that is complementary to atom-coordinate-based 3D molecular representation learning. The task-wise heatmaps include ESOL, FreeSolv, Lipophilicity, BBBP, BACE, ClinTox, and SIDER.

**Figure S2.**
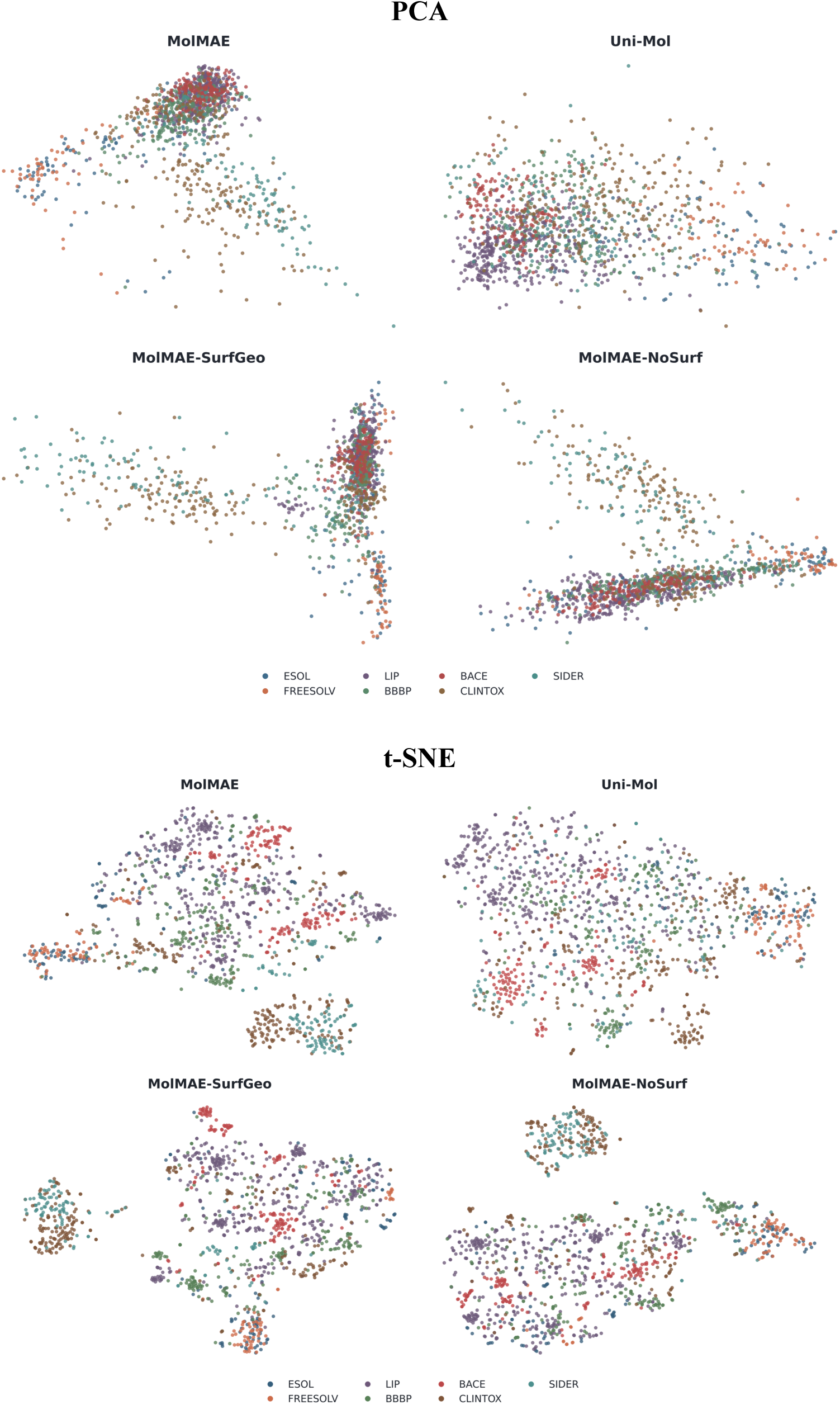
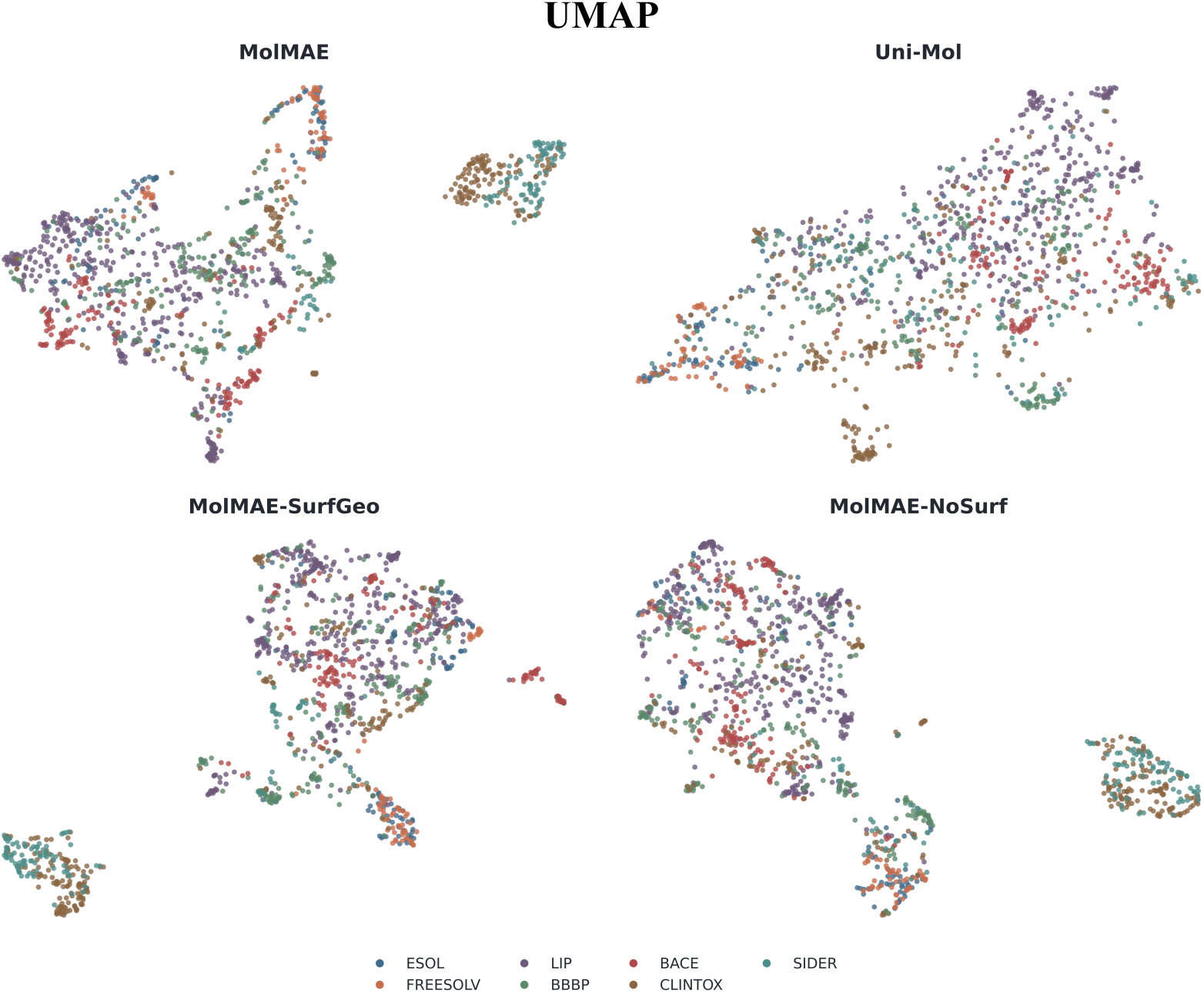
Chemical-space visualization of MolMAE, Uni-Mol, and surface ablation variants. Low-dimensional projections of molecular embeddings are shown for MolMAE, Uni-Mol, MolMAE-SurfGeo, and MolMAE-NoSurf across the seven downstream datasets. Each point represents one molecule, and colors denote dataset membership. The visualization illustrates that the four encoders organize the same molecular collections differently, consistent with the quantitative representation-similarity results. These differences suggest that molecular surface information changes the learned chemical-space geometry rather than simply reproducing the embedding structure of Uni-Mol.

**Figure S3.**
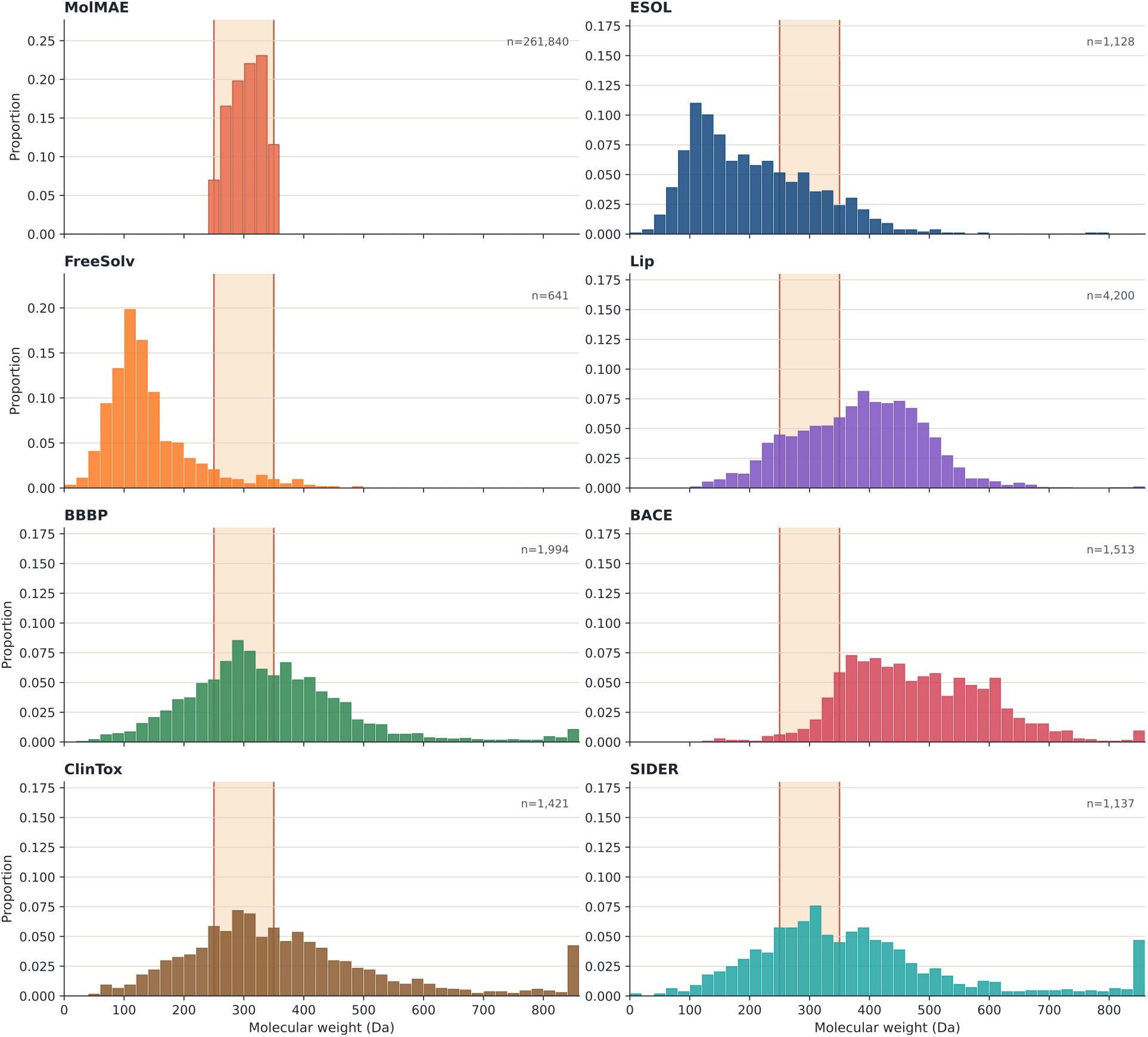
Molecular-weight distributions of the MolMAE pretraining corpus and downstream benchmarks.

The molecular-weight distribution of the MolMAE pretraining corpus is compared with the seven downstream benchmarks using 20 Da bins. The shaded region marks the lead-focused molecular-weight window. The pretraining corpus is concentrated in a lead-like regime, whereas the downstream datasets exhibit different molecular-weight profiles. This analysis highlights possible distributional differences between pretraining and downstream evaluation and provides context for task-dependent transfer performance.

**Figure S4.**
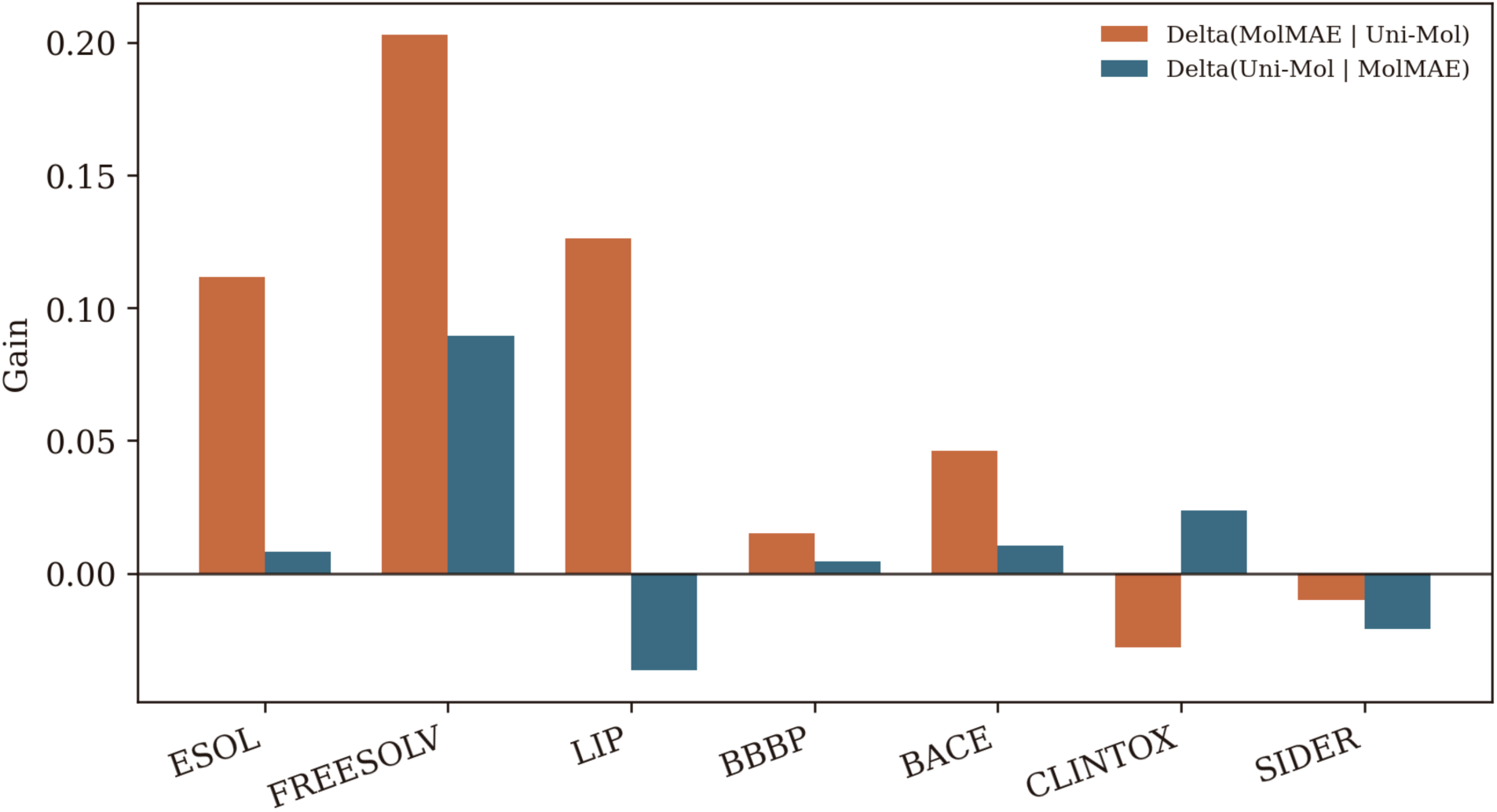
Complementary predictive gains from combining MolMAE and Uni-Mol embeddings. The bar plot reports the change in linear-probe performance when MolMAE and Uni-Mol embeddings are concatenated and used jointly, compared with single-model embeddings. Positive gains indicate that the two representations contain complementary predictive information. This analysis further tests whether MolMAE captures non-redundant surface-guided features beyond Uni-Mol’s atom-coordinate-based 3D representation.

**Table S1.** ESOL comparison with additional supervised baselines. Results are reported under the same scaffold-split evaluation protocol used in this study. Lower RMSE indicates better performance. Chemprop is reported with the best hyperparameter setting selected from the grid-search protocol, while Uni-Mol uses the official recommended setting. Bold indicates the best result.

| Model | RMSE |
| --- | --- |
| MolMAE | **0.740** |
| AttentiveFP | 0.760 |
| Chemprop | <u>0.747</u> |
| Uni-Mol | 0.866 |

